# Application of probiotic *Bacillus* spp. isolated from African nightcrawler (*Eudrilus eugeniae*) on Nile Tilapia (*Oreochromis niloticus* L.)

**DOI:** 10.1101/2020.03.08.982819

**Authors:** J. Samson, K.M. Quiazon, C. Choresca

## Abstract

Due to the emergence of antibiotic-resistant pathogens, probiotics in aquaculture are used for the prevention of infectious microbial diseases and substitute for antibiotics and chemotherapeutics. In this study, we evaluated the effect of probiotic *Bacillus* spp. isolated from African nightcrawler (*Eudrilus eugeniae*) on the growth, feed utilization, and disease resistance of Nile tilapia (*Oreochromis niloticus*). Four probiotic strains of *Bacillus* spp. (ANSCI9, BFAR9, RM3, and RM10) were individually incorporated in the commercial diet (control) at 10^8^ CFU g^-1^ of feed. The experimental fish were fed at 5% of their body weight for 30 days, and subjected to a 14-day *Aeromonas hydrophila* challenge test afterward. The results showed the probiotic-treated groups have higher (*P*<0.05) average body weight (ABW) (4.51 ± 0.34 g) than the control (3.89 ± 0.17 g). The BFAR9 (2.73 ± 0.26 g) and RM10 (3.15 ± 0.30 g) showed higher (*P*<0.05) absolute growth (AG) than the control (2.20 ± 0.16 g). Furthermore, RM10 had higher (*P*<0.05) specific growth rate (SGR) (1.60 ± 0.10 % day^-1^) and relative growth rate (RGR) (181.39 ± 18.16 %) than the control (SGR=1.29 ± 0.07 % day^-1^; RGR=129.84 ± 9.77 %). Consequently, RM10 had significantly lower (*P*<0.05) feed conversion ratio (FCR) (1.99 ± 0.13) than the control (2.60 ± 0.16). The challenge test revealed that the probiotic-treated groups have higher (*P*<0.05) survival (81.25 ± 9.57 %) than the control (55.00 ± 19.15 %). These results revealed that the probiotic *Bacillus* spp. isolated from *E. eugeniae* improved the growth, feed utilization, and the disease resistance of Nile tilapia.

## Introduction

The need for sustainable aquaculture has encouraged exploration into the use of feed additives for growth promotion and health improvement of aquatic organisms (Zokaeifar et al. 2012; Saleh et al. 2015; Michael et al. 2017; Samson 2019b, a). These feed additives such as the probiotic bacteria are used for the growth promotion, pathogen inhibition, enhanced nutrient utilization, water remediation, stress tolerance, and reproduction of aquatic species (Balcazar et al. 2006; Cruz et al. 2012). These beneficial microorganisms can be isolated from various sources, including in food (Leite et al. 2015; Demir and Başbülbül 2017), aquatic organisms (Lin et al. 2013; Lamari et al. 2014; Sanchez-Ortiz et al. 2015), terrestrial animals (Brashears et al. 2003; Musikasang et al. 2009; Jena et al. 2013; Feng et al. 2017), and humans (Amenu 2015; Reis et al. 2016). In this study, we evaluated the probiotic potential of the isolated *Bacillus* spp. from earthworms on Nile tilapia. Although some studies have used earthworms as a protein source for fish (Boaru et al. 2016; Mohanta et al. 2016), and explored the presence of microorganisms in them (Horn et al. 2003; Singleton et al. 2003; Kim et al. 2004; Byzov et al. 2009), to the best of our knowledge, no study has demonstrated the effect of these microorganisms on aquatic species.

The gut of earthworms provides an ideal and favorable environment for the growth and activity of bacteria (Govindarajan and Prabaharan 2015). For several centuries, China and other parts of the Far East believed that earthworms promote overall health, and are sources of therapeutic drugs for various diseases (El-Kamali 2000; Ismail 2005). Furthermore, indigenous people all over the world, more particularly in Asia, including Vietnam, China, Korea, India, and Myanmar, extract and use the biologically active compounds from earthworms (Ranganathan 2006). Unfortunately, there is no available information on the application of probiotic bacteria from earthworms, specifically on aquaculture. Hence, in this present study, we aimed to evaluate the effect of probiotic *Bacillus* spp. isolated from *E. eugeniae* on the growth, feed utilization, and disease resistance of Nile tilapia.

## Materials and Methods

### Source of Bacteria

Four probiotic strains of *Bacillus* spp. (ANSCI9, BFAR9, RM3, and RM10) previously isolated from *E. eugeniae* were used in this study (Samson et al. 2020). The 16s rRNA sequences of the probiotic isolates are available in the GenBank, with accession numbers: MH919310, MH919302, MH919306, and MH919308.

### Experimental Diets and Design

A commercial diet (Tateh Aquafeeds, Philippines) served as the basal diet (control). The proximate composition was as follows: crude protein 31%, lipid 5%, crude fiber 8%, and crude ash 12%. The probiotic-treated diets were prepared to contain a single strain of *Bacillus* spp. (ANSCI9, BFAR9, RM3, and RM10) per group. The McFarland standard was used to estimate the bacterial densities. Dilutions were done using phosphate buffer solution (PBS) to have a similar bacterial density. The isolates were sprayed on commercial feeds (10^8^ CFU·g^-1^) and air-dried for 2 hours before storing at 4°C.

Four hundred *O. niloticus* fingerlings were distributed into 20 glass aquaria (52 × 25 × 30 cm) divided into five experimental groups (four replicates per group). Each group consists of 80 fish (20 fish × 4 aquaria), which had an initial weight of 1.76 ± 0.07 g. The water quality parameters during the study were the following: the temperature at 25.3 ± 0.9 °C, pH at 8.0 ± 0.2, and dissolved oxygen at 3.52 ± 1.16 ppm. The fish were fed their respective diets (at approximately 5% of body weight day^-1^) thrice daily at 8:00, 12:00, and 16:00 for 30 days.

### Data Collection

At the end of the feeding trial, all fish were measured and weighed. Growth parameters and survival rate were measured as follows:

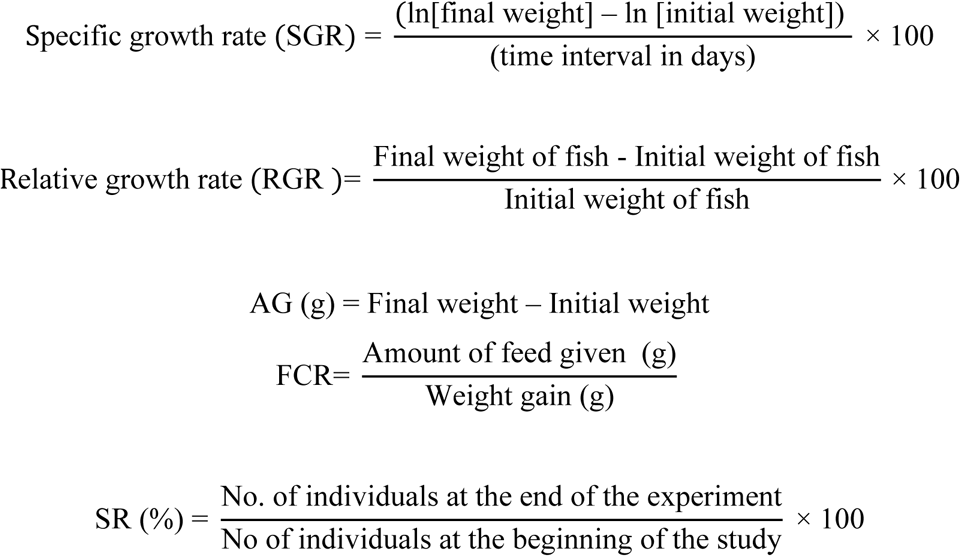

### Challenge Test

*O. niloticus* fingerlings (4.39 ± 0.40 g) were challenged with *A. hydrophila* (BIOTECH 10089) after the 30-day feeding trial. Fish were anesthetized in tricaine methanesulfonate (MS222) solution and intramuscularly injected with 0.1 mL *A. hydrophila* suspension (10^7^ CFU·mL^-1^). The mortality was observed daily until the 14^th^ day, and the relative level of protection (RLP) among the challenged fish was determined using the equation: [100 – (immunized mortality% / control mortality%)] × 100 (Ruangroupan et al. 1986)

### Statistical Analysis

Results were subjected to a one-way analysis of variance (ANOVA), followed by Duncan’s Multiple Range Test (DMRT) at a significant level of *P<*0.05 when there was a significant difference among the treatments.

## Results

### Growth Performance

The summary of the measurements and growth parameters of the experimental fish is shown in Table 1. The treatments supplemented with probiotic *Bacillus* spp. showed higher (*P*<0.05) ABW than the control. RM10 gained the highest ABW (4.88 ± 0.30 g) among the treatments; whereas, the lowest was recorded from the control group (3.89 ± 0.17 g). Treatments RM10 (6.91 ± 0.23 cm), BFAR9 (6.88 ± 0.07 cm), and RM3 (6.65 ± 0.19 cm) have significantly higher (*P*<0.05) lengths than the control (6.16 ± 0.11 cm). Also, treatment RM10 had significantly higher (*P*<0.05) SGR (1.60 ± 0.10 % day^-1^) and RGR (181.39 ± 18.16 %) than the control (SGR=1.29 ± 0.07 % day^-1^; RGR=129.84 ± 9.77 %). In addition, treatments BFAR9 (2.73 ± 0.26 g) and RM10 (3.15 ± 0.30 g) showed significantly higher (*P*<0.05) AG than the control (2.20 ± 0.16 g). Consequently, treatment RM10 had significantly lower (*P*<0.05) FCR (1.99 ± 0.13) than the control (2.60 ± 0.16). While no significant difference was observed on the survival of the treatments.

**Table 1.** Summary (Mean ± SD) of measurements and growth parameters of the experimental fish (n=4).

|  | TREATMENTS |  |  |  |  |
| --- | --- | --- | --- | --- | --- |
|  | Control | ANSCI9 | BFAR9 | RM3 | RM10 |
| Initial ABW (g) | 1.69 ± 0.04<sup>a</sup> | 1.80 ± 0.07<sup>a</sup> | 1.78 ± 0.09<sup>a</sup> | 1.77 ± 0.08<sup>a</sup> | 1.74 ± 0.02<sup>a</sup> |
| Initial Length (cm) | 4.63 ± 0.21<sup>a</sup> | 4.59 ± 0.12<sup>a</sup> | 4.82 ± 0.24<sup>a</sup> | 4.67 ± 0.10<sup>a</sup> | 4.66 ± 0.08<sup>a</sup> |
| Final ABW (g) | 3.89 ± 0.17<sup>c</sup> | 4.31 ± 0.20<sup>b</sup> | 4.51 ± 0.21<sup>ab</sup> | 4.34 ± 0.35<sup>b</sup> | 4.88 ± 0.30<sup>a</sup> |
| Final Length (cm) | 6.16 ± 0.11<sup>c</sup> | 6.18 ± 0.04<sup>c</sup> | 6.88 ± 0.07<sup>a</sup> | 6.65 ± 0.19<sup>b</sup> | 6.91 ± 0.23<sup>a</sup> |
| AG (g) | 2.20 ± 0.16<sup>c</sup> | 2.51 ± 0.23<sup>bc</sup> | 2.73 ± 0.26<sup>b</sup> | 2.57 ± 0.35<sup>bc</sup> | 3.15 ± 0.30<sup>a</sup> |
| SGR (% day<sup>-1</sup>) | 1.29 ± 0.07<sup>b</sup> | 1.35 ± 0.11<sup>b</sup> | 1.44 ± 0.13<sup>ab</sup> | 1.39 ± 0.14<sup>b</sup> | 1.60 ± 0.1<sup>a</sup> |
| RGR (%) | 129.84 ± 9.77<sup>b</sup> | 139.71 ± 16.82<sup>b</sup> | 153.16 ± 21.1<sup>ab</sup> | 145.61 ± 22.44<sup>b</sup> | 181.39 ± 18.16<sup>a</sup> |
| FCR | 2.60 ± 0.16<sup>a</sup> | 2.47 ± 0.23<sup>a</sup> | 2.30 ± 0.21<sup>ab</sup> | 2.41 ± 0.31<sup>ab</sup> | 1.99 ± 0.13<sup>b</sup> |
| SR (%) | 95.00 ± 5.77<sup>a</sup> | 87.50 ± 25.00<sup>a</sup> | 97.50 ± 2.89<sup>a</sup> | 97.50 ± 5.00<sup>a</sup> | 91.25 ± 14.36<sup>a</sup> |

Note: Treatments with the same superscript are not significantly different at $P < 0.05$ .

### Challenge Test

The cumulative mortality of the experimental groups was recorded daily until the 14^th^ day (Fig 1). All treated groups showed significantly higher (*P*<0.05) survival rates rate (81.25 ± 9.57 %) than the control (55.00 ± 19.15 %). Treatment RM10 had the highest survival rate (85.0 ± 10.00 %) among the treatments, while the lowest was recorded on the control (55 ± 19.15%). The RLP of the treated groups is ≥50% with the highest value in RM10 (66.67%) while the lowest for RM3 (50.00%). The result of the challenge test is presented in Table 3.

**Table 3.** Values (Mean ± SD) of SR RLP for each treatment after challenging with *A. hydrophila.*

|  | TREATMENTS |  |  |  |  |
| --- | --- | --- | --- | --- | --- |
|  | Control | ANSCI9 | BFAR9 | RM3 | RM10 |
| SR (%) | 55.0 ± 19.15<sup>b</sup> | 82.5 ± 8.16<sup>a</sup> | 80.0 ± 10.00<sup>a</sup> | 77.5 ± 20.82<sup>a</sup> | 85.0 ± 10.00<sup>a</sup> |
| RLP (%) | 0.0 | 61.11 | 55.56 | 50.00 | 66.67 |

Note: Treatments with the same superscript are not significantly different at $P < 0.05$ .

**Figure 1.**
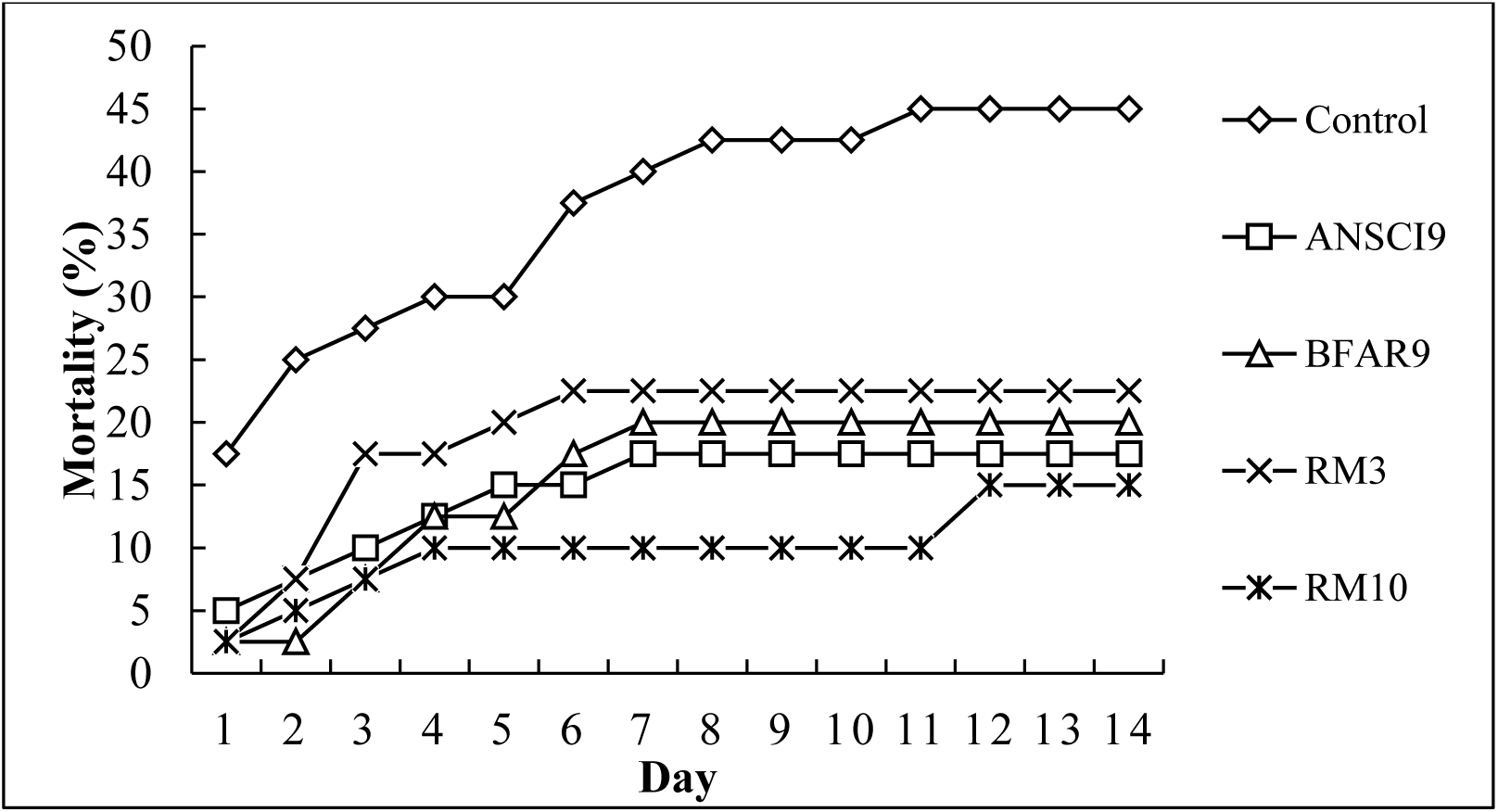
Cumulative mortality of each treatment during the *A. hydrophila* challenge test.

## Discussion

The beneficial effect of probiotic *Bacillus* spp. isolated from *E. eugeniae* was observed in this study. Nile tilapia showed better growth when fed with probiotics. Previous studies have reported similar results on the effect of *Bacillus* spp. on the growth and survival of Nile tilapia. Soltan et al., (2016) reported that commercial probiotics containing allicin, high unit hydrolytic enzyme, *Bacillus subtilis* spores, and ginseng extract improved the growth and feed utilization of Nile tilapia. Similarly, *Bacillus pumilus* improved the growth performance of Nile tilapia due to enhanced immune system, health status, and disease resistance (Aly et al. 2008). Furthermore, the application of *Bacillus coagulans* and other *Bacillus* spp. in the water also enhanced the growth and immune response of Nile tilapia (Zhou et al. 2009; Sutthi et al. 2018). According to Buruian et al. (2014), the application of probiotic *Bacillus* spp. in the aquaculture system improves the growth performance, immune response, disease resistance, and survival of the fish. These probiotics produce extracellular enzymes (proteases, amylases, and lipases), and provide vitamins, fatty acids, and amino acids that have a beneficial effect on the metabolism of aquatic animals (Balcazar et al. 2006). The observed growth improvement of treated groups can be a result of improved enzymatic activity on the fish gut, as the probiotics used in the experiment produce extracellular enzymes. These extracellular enzymes aids in breaking down food molecules, which makes the absorption of nutrients more efficient (EL-Haroun et al. 2006). In contrast, Silva et al. (2015) reported that the addition of *Bacillus amyloliquefaciens* did not improve the growth and proximal composition of Nile tilapia; however, they observed a significant increase of villi height and number of goblet cells in the intestine. Their result suggests that the digestion and nutrient absorption of fish had been improved. Similarly, Shelby et al. (2006) observed no significant improvement in growth; but they recorded higher survival rates after 39 to 63 days of probiotic application. As we noted in this study, only RM10 showed significantly better SGR, RGR, and FCR; although the probiotics used are of the same genus (i.e., *Bacillus*) and source (i.e., *E. eugeniae*). This result does not imply that the three probiotic isolates (i.e., ANSCI9, BFAR9, RM3) are less efficient. In fact, all probiotic-treated groups showed higher survival rates when challenged with *A. hydrophila*. This suggests that the effect of the probiotics may not be observed on growth but rather to the immunity and intestinal morphology of the fish. Thus, it is better to evaluate the effect of these probiotics in the immunological parameters and intestinal morphology of the fish. Furthermore, this observation also suggests that the effects of probiotics application differs from each bacterial species and strain. Correspondingly, it is important to measure the effective rate of application for each probiotic bacterium.

In this study, higher survival rates of probiotic-treated groups infected with *A. hydrophila* was recorded. Similar results were reported by Selim and Reda (2015); they reported that dietary supplementation of *B. amyloliquefaciens* improved the survival of Nile tilapia infected with *Yersinia ruckeri* or *Clostridium perfringens* type D. Moreover, Gupta et al. (2014) also observed an improvement in the survival of *Cyprinus carpio* fry fed with probiotics and challenged with *A. hydrophila.* In our study, the improvement of survival rates of the probiotic-treated groups is attributed to the ability of probiotics to enhance the immune system. It is a result of immunity modulation or immunomodulation, which is one of the common health benefits credited to the application of probiotics (Cross 2002). It is well-known that probiotics may lessen the occurrence of disease or decrease the danger of disease outbursts. One possible mechanism for this is the prevention of pathogen colonization on the intestinal mucosa or other epithelial surfaces; as probiotic strains colonize these surfaces (Balcazar et al. 2006). Furthermore, they can also produce inhibitory substances against pathogenic organisms and compete for essential nutrients (Ringø and Gatesoupe 1998). The spore-forming ability of the *Bacillus* spp. used in this study may perhaps aid in administering their effect. This ability increases their chance to survive the harsh conditions in the intestine and the activities of the digestive enzymes. To further explore the beneficial effects of these probiotic bacteria, we suggest assessing their effects on the biochemical and hematological parameters of the fish. Additionally, the ability of these probiotics in preventing the infection of other pathogenic bacteria should also be considered.

## Conclusion

This study is the first report on the application of probiotic *Bacillus* spp. isolated from *E. eugeniae* in fish. The results of this study revealed the beneficial effect of *Bacillus* spp. isolated from *E. eugeniae* on the growth performance, feed utilization and disease resistance of Nile tilapia.

## Acknowledgement

The authors acknowledge the financial support from the Department of Science and Technology -Accelerated Science and Technology Human Resource Development Program (DOST-ASTHRDP). Also, the authors acknowledge the assistance from the Fisheries Biotechnology Center, College of Fisheries, and Freshwater Aquaculture Center, CLSU.

